# Network reconstruction and systems analysis of plant cell wall deconstruction by *neurospora crassa*

**DOI:** 10.1101/133165

**Authors:** Areejit Samal, James P. Craig, Samuel T. Coradetti, J. Philipp Benz, James A. Eddy, Nathan D. Price, N. Louise Glass

## Abstract

Plant biomass degradation by fungal derived enzymes is rapidly expanding in economic importance as a clean and efficient source for biofuels. The ability to rationally engineer filamentous fungi would facilitate biotechnological applications for degradation of plant cell wall polysaccharides. However, incomplete knowledge of biomolecular networks responsible for plant cell wall deconstruction impedes experimental efforts in this direction. To expand this knowledge base, a detailed network of reactions important for deconstruction of plant cell wall polysaccharides into simple sugars was constructed for the filamentous fungus *Neurospora crassa*. To reconstruct this network, information was integrated from five heterogeneous data types: functional genomics, transcriptomics, proteomics, genetics, and biochemical characterizations. The combined information was encapsulated into a feature matrix and the evidence weighed to assign annotation confidence scores for each gene within the network. Comparative analyses of RNA-seq and ChIP-seq data shed light on the regulation of the plant cell wall degradation network (PCWDN), leading to a novel hypothesis for degradation of the hemicellulose mannan. The transcription factor CLR-2 was subsequently experimentally shown to play a key role in the mannan degradation pathway of *Neurospora crassa*. Our network serves as a scaffold for integration of diverse experimental data, leading to elucidation of regulatory design principles for plant cell wall deconstruction by filamentous fungi, and guiding efforts to rationally engineer industrially relevant hyper-production strains.

## Introduction

Plant biomass, primarily composed of lignocellulose, is a renewable and environmentally-clean energy source, and a promising feedstock for the production of next generation biofuels and specialty chemicals (1-3). A principal barrier for economical production of biofuels is the high production cost of biomass depolymerization enzymes (4). Filamentous fungi are among the most efficient degraders of lignocellulosic biomass in nature and play a key role in carbon recycling (5,6). Industrially relevant strains, such as *Trichoderma reesei* were constructed through multiple rounds of random mutagenesis and can secrete up to 100 g/L of protein (7,8). However, rationally engineering strains of filamentous fungi to further enhance secretion of enzymes is a major challenge in bioenergy research (9). To meet this challenge and aid future experimental efforts, a system-level understanding of plant cell wall deconstruction by filamentous fungi is necessary (5,10).

The model filamentous fungus *Neurospora crassa* has well-developed genetics, biochemistry, molecular biology and a well annotated genome (11-14). In nature, *N. crassa* colonizes freshly burnt plant biomass and shows robust growth on lignocellulose (5,15-19). The suite of experimental resources available for *N. crassa* makes it an ideal model system for bioenergy-related research, particularly for the elucidation of plant cell wall deconstruction mechanisms (5,16-20). Research on *N. crassa* contributed to the discovery of a new class of enzymes called lytic polysaccharide monooxygenases (LPMOs), which greatly increase synergy in cellulose degradation. In addition, novel cellodextrin transporters from *N. crassa* were utilized to engineer improved yeast strains for sugar fermentation (21-24). A network reconstruction encompassing the present knowledge of metabolic reactions, enzymes, and associated genes in *N. crassa* dedicated to the deconstruction of plant cell wall polysaccharides into simple fermentable sugars will further expedite experimental efforts.

The availability of fully sequenced genomes and accumulated wealth of biochemical evidence led to the reconstruction of genome-scale and manually curated metabolic networks for more than 50 organisms across the three domains of life (25,26). These genome-scale metabolic networks have been widely analyzed using constraint-based modeling methods to predict the response to environmental and genetic perturbations (27,28). Notably, only a few curated genome-scale metabolic reconstructions have been built for filamentous fungi (29-34). While a manually curated genome-scale metabolic network for *N. crassa* exists (34), this reconstruction and those built for other ascomycete fungi (29-34) are limited by significant knowledge gaps, specifically pathways for the degradation and utilization of plant cell wall polysaccharides.

To overcome this limitation, we built a detailed network of biochemical reactions important for the degradation of plant cell wall polysaccharides into simple fermentable sugars in *N. crassa* (Figure 1; Table S1). Plant cell walls are largely composed of complex polysaccharides that include cellulose, hemicellulose and pectin (5,35-40) (Figure 1; Table S1). Cellulose is the most abundant plant cell wall polysaccharide and is an unbranched structure composed of linear chains of β-1,4-linked D-glucose residues. The second most abundant is the heterogeneous group of hemicelluloses, composed of several branched polymers, including xylan, xyloglucan, mannan, and mixed-linkage glucan. Pectin is a minor constituent of mature plant cell walls but the most complex heteropolysaccharide. Its main constituents are homogalacturonan, xylogalacturonan, and rhamnogalacturonan I.

**Figure 1:**
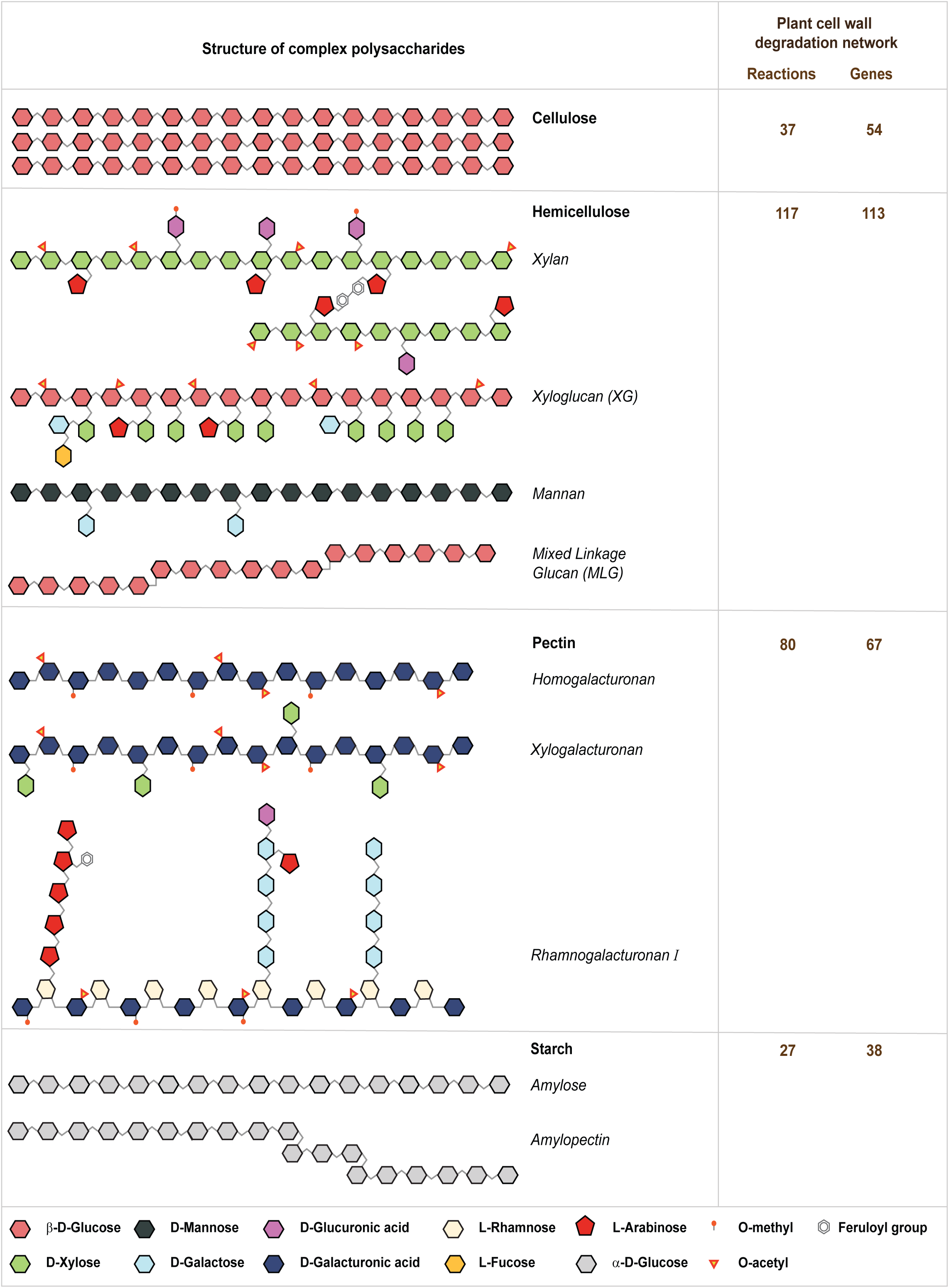
Schematic illustration of the structure of different plant cell wall polysaccharides along with the associated reactions and genes in the PCWDN of *N. crassa*. Cellulose has an unbranched structure composed of linear chains of β-1,4-linked D-glucose residues. Hemicellulose comprises several branched polymers including xylan, xyloglucan (XG), mannan and mixed-linkage glucan (MLG). Pectin is a family of several polymers including homogalacturonan, xylogalacturonan and rhamnogalacturonan I. Starch is a polymer composed of amylose and amylopectin. On the right, the number of reactions and genes involved in the degradation of cellulose, hemicelluloses, pectin and starch are indicated that are captured in our PCWDN.

The plant cell wall degradation network (PCWDN) reconstruction and annotation pipeline described here involved the integration of five heterogeneous data types: functional genomics, transcriptomics, proteomics, genetics, and biochemical information, along with extensive manual curation based on more than 130 research articles (Figure 2; Table S1). The combined annotation information was encapsulated in a feature matrix, which was used to assign annotation confidence scores to PCWDN genes. Comparative analysis of RNA sequencing (RNA-seq) based global transcriptome profiles underlined the importance of PCWDN genes for adaptations to different plant cell wall polysaccharides. Subsequent analyses of RNA-seq and ChIP-seq data within the context of the *N. crassa* PCWDN led to novel insights on the roles of key transcription factors (TFs) in the deconstruction of plant biomass, which were tested here. The *N. crassa* PCWDN will serve as a scaffold for the integration and systems analyses of diverse experimental data, helping to elucidate the regulatory design principles underlying plant cell wall deconstruction by filamentous fungi.

**Figure 2:**
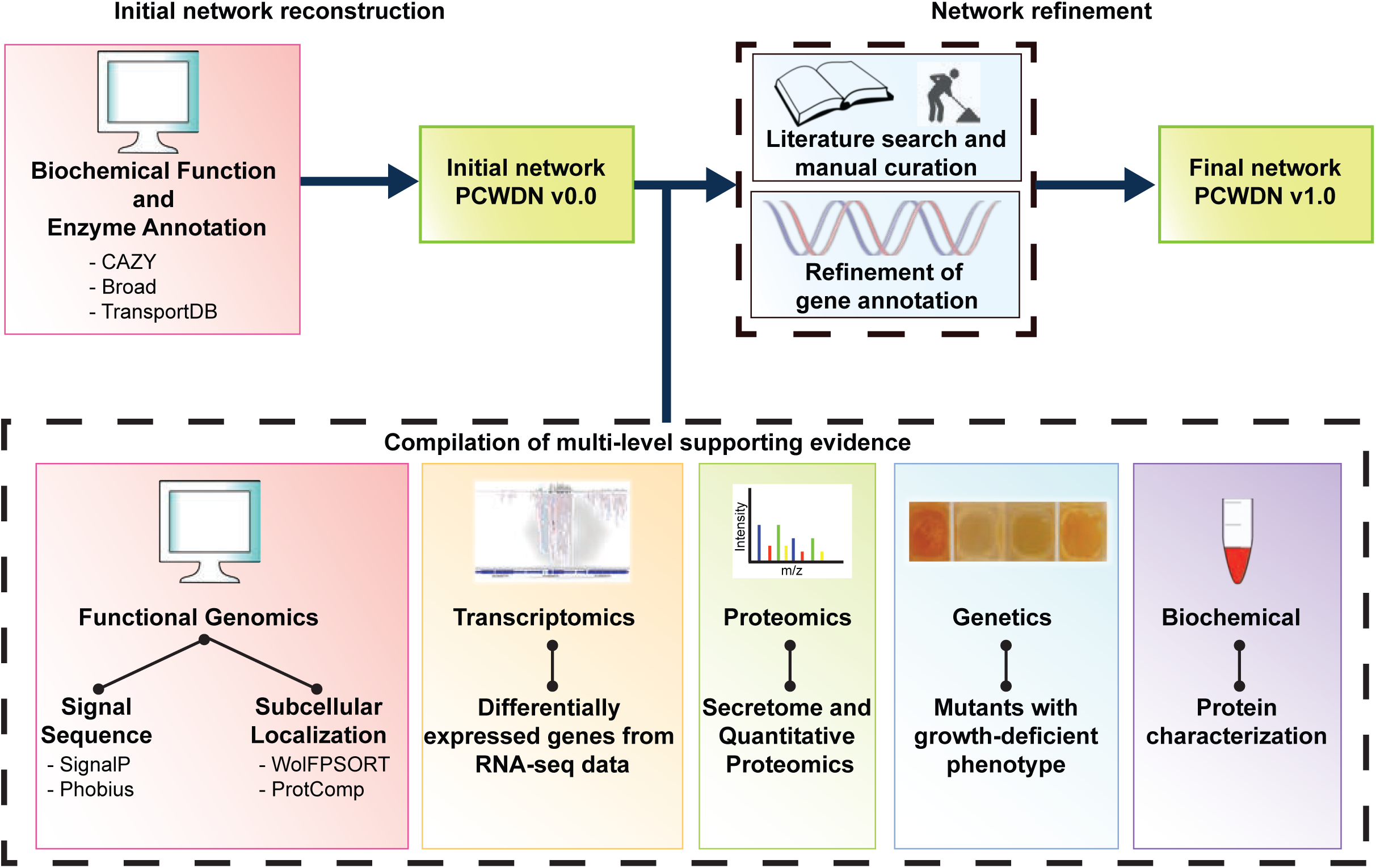
Schematic illustration of the pipeline for reconstruction and annotation of the PCWDN of *N. crassa*. An initial scaffold network, PCWDN v0.0, was assembled based on annotation information in several databases. Extensive literature-based manual curation was then performed to fill knowledge gaps in the initial PCWDN v0.0. Gene annotations in the final PCWDN v1.0 were refined based on multi-level supporting evidence from five heterogeneous data types.

## Results and Discussion

### Network Reconstruction and Annotation Pipeline

We assembled an initial list of biochemical reactions and associated genes in the PCWDN v0.0 of *N. crassa* by combining information on predicted enzymes and transporters involved in degradation of plant cell wall polysaccharides from the following sources: the Carbohydrate-Active enZYmes database (CAZY) (41), the *N. crassa* e-Compendium (42), the genome annotation for *N. crassa* OR74A (12,13), and TransportDB (43) (Methods; Figure 2). Specifically, 110 out of 231 CAZY genes predicted to encode carbohydrate-active enzymes in the genome were included in the PCWDN v0.0 (Table S2). The remaining 121 CAZY genes mainly belong to families of enzymes active on chitin and chitosan and thus are not likely to be involved in the PCWDN, but rather in remodeling the fungal cell wall (Table S2).

The annotation of PCWDN genes in the above databases has not been updated with data on plant cell wall deconstruction by *N. crassa*. For example, the current OR74A genome annotation is unable to differentiate between cellulolytic LPMOs, hemicellulose-active LPMOs, and starch-active LPMOs (44-50). Thus, we performed extensive literature-based manual curation involving more than 130 research articles (Table S1) to fill the knowledge gaps in the initial PCWDN v0.0 and compiled multi-level supporting evidence as described below from five heterogeneous data types: functional genomics, transcriptomics, proteomics, genetics and biochemical characterizations, to annotate genes in the final PCWDN v1.0 of 202 reactions and 168 genes (Figure 2; Table S1). The 202 reactions in the final PCWDN of *N. crassa* were further subdivided into 101 extracellular reactions, 35 transport reactions, and 66 intracellular reactions (Table S1).

#### Functional genomics-based annotation

An important annotation feature of PCWDN enzymes is their predicted subcellular localization. For example, the hydrolysis of cellodextrins into D-glucose by β-glucosidases can occur in the extracellular space or in the intracellular space (Table S1). Of the 168 PCWDN genes, products of 103, 19 and 46 genes are associated with extracellular, transport and intracellular reactions, respectively. We used SignalP (51) and Phobius (52) to predict the presence of signal peptides in PCWDN proteins to determine if they were destined towards the secretory pathway (Methods). We found that 89 out of the 103 gene products (∼86%) associated with extracellular reactions were predicted to have a signal peptide by at least one of the two tools, while no gene products associated with transport or intracellular reactions were predicted to have a signal peptide by either of the two tools (Table S3). WoLF PSORT (53) and ProtComp was also used to predict subcellular localization of proteins (Methods). Predictions from at least one of the two tools matched the assigned localization for 90 out of the 103 gene products (∼87%) associated with extracellular reactions, while the predictions from at least one of the two tools matched the assigned localization for all gene products associated with transport or intracellular reactions in the PCWDN (Table S3).

Using the compiled functional genomics-based information (Table S3), we assessed the annotation support for the 168 PCWDN members. Specifically, functional genomics information was considered to support the annotation of a PCWDN enzyme if the following three conditions were satisfied (Figure 3): (i) Gene annotation in CAZY database (41) or Broad OR74A genome (12,13) or TransportDB (43) matches the assigned biochemical function in the network; (ii) SignalP or Phobius predicts the presence of a signal peptide in extracellular (secreted) enzymes and the absence of a signal peptide in intracellular enzymes; (iii) Subcellular localization predictions from WoLF PSORT or ProtComp matches the assigned localization in the network. Based on this definition, we obtained functional genomics-based annotation support for 145 of the 168 PCWDN genes (Table S3). *Transcriptomics-based annotation*

**Figure 3:**
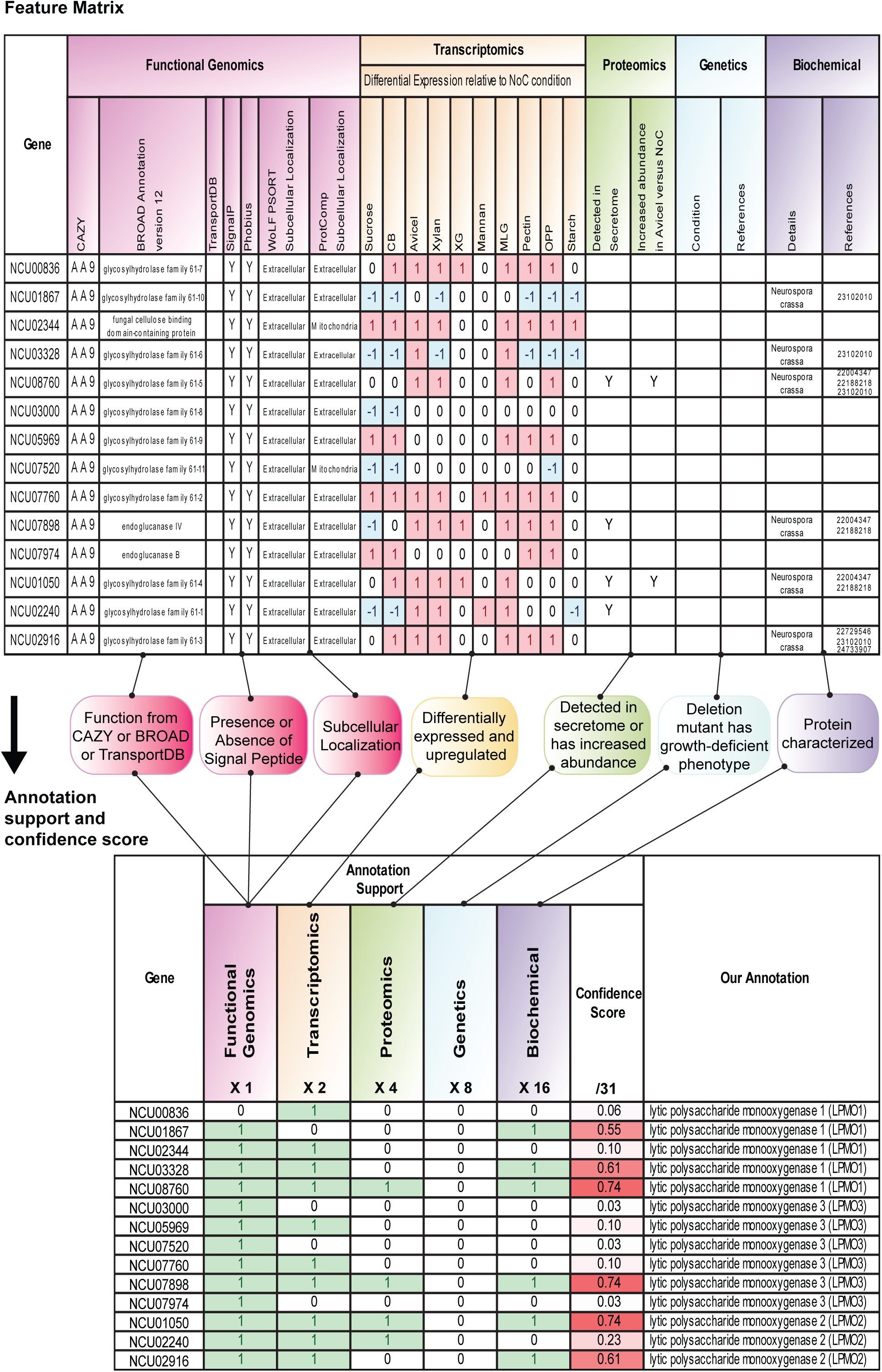
Feature matrix and annotation confidence scores for genes encoding AA9 LPMOs in *N. crassa*. Combined annotation information from the five different data types was captured in a feature matrix, and a method was devised to assign annotation confidence scores to PCWDN genes. A differential weighting system was used to account for the different levels of confidence associated with the information from each data type. The majority of genes encoding AA9 LPMOs of class 1 (3 out of 5 genes) and class 2 (2 out of 3 genes) are well characterized, while only 1 out of 6 genes encoding AA9 LPMOs of class 3 is well characterized. NoC: No Carbon; CB: Cellobiose; XG: Xyloglucan; MLG: Mixed-Linkage Glucan, OPP: Orange Peel Powder.

To augment the annotation of PCWDN genes, we used RNA-seq data and compared transcriptional profiles of *N. crassa* WT strain (FGSC 2489 (12); Methods) grown under different carbon source regimes corresponding to the different carbohydrates that make up the plant cell wall. Previous studies (16,19) generated RNA-seq data from shift experiments, in which a 16 hr-old culture of *N. crassa* WT was shifted for 4 hrs to minimal media with no carbon (NoC) source or one of five carbon sources: sucrose, cellobiose (CB), Avicel (microcrystalline cellulose), xylan, pectin or orange peel powder (OPP, a pectin rich substrate) (Methods; Table S4). We replicated this experimental design, generating RNA-seq data for four additional carbon sources: xyloglucan (XG), mannan, mixed-linkage glucan (MLG), and starch (Methods; Table S4). This approach ensured that a comparative analysis of transcriptional profiles could be performed between all tested plant cell wall polysaccharide. A pipeline consisting of standard software was used to analyze the RNA-seq data and identify differentially expressed genes between the nine different conditions corresponding to different polysaccharides and the two controls corresponding to NoC and sucrose conditions (Methods; Table S4).

Based on the compiled RNA-seq data, we generated transcriptomics-based annotation support for the PCWDN genes (Figure 3; Table S4). We considered RNA-seq data to support the annotation of a gene involved in the degradation of a specific polysaccharide if its transcript was significantly differentially expressed and upregulated in the relevant carbon source as compared to the NoC control. For example, RNA-seq data supported the annotation of the gene *gh6-3* (NCU07190), predicted to encode an exo-β-1,4-glucanase (cellobiohydrolase) involved in cellulose degradation, as this gene is upregulated on Avicel in comparison to the NoC control (Table S4). We obtained transcriptomics-based annotation support for 106 of the 168 PCWDN genes (Table S4).

#### Proteomics-based annotation

To further enrich the PCWDN and provide an added layer of confidence in the annotation, we compiled proteomics data from previous *N. crassa* studies (15,17-19,54-57) (Table S5). In these studies, the secretomes from *N. crassa* grown on sucrose, Avicel, xylan, pectin, OPP, and NoC had been characterized using a shotgun proteomics approach (15,17-19,54,55) or by quantitative proteomics (57) (Table S5). The compiled data were used to generate proteomics-based support for the annotation of PCWDN genes. Proteomics data supported the annotation of an enzyme if the protein was detected in the secretome or displayed an increased abundance in a carbon source as compared to the NoC control (Figure 3). For example, proteomics data further supported the annotation of *gh6-3*, as the encoded protein was detected in the secretome and also increased in abundance on Avicel compared to the NoC control (Table S5). In this way, we obtained proteomics-based annotation support for 68 of the 168 PCWDN genes (Table S5). In cases where an enzyme in the PCWDN was detected in a secretome but was not predicted to be secreted using functional genomics tools, the proteomics-based evidence was considered more reliable and given priority. This approach led to the re-annotation of four PCWDN genes: *gla-1*(NCU01517), *gh6-3* (NCU07190), *gh5-7* (NCU08412) and *ce5-2* (NCU09663).

#### Genetics-based annotation

Next, we mined published literature to compile an experimentally verified dataset of *N. crassa* deletion strains for PCWDN genes with a growth-deficient phenotype as compared to the parental WT strain (FGSC 2489) (Table S6). This dataset was used to assign genetics-based annotation support to PCWDN enzymes. For example, the genetics-based dataset supported the annotation of the gene *gh10-2* (NCU08189) as an endo-β-1,4-xylanase involved in xylan degradation, since the deletion strain for this gene exhibited a growth-deficient phenotype on xylan (17) (Table S6). Overall, genetics-based information supported the annotation of 19 out of the 168 PCWDN genes (Table S6).

#### Biochemical characterization of enzymes

Lastly, we mined the literature for biochemical data to support the annotation of PCWDN enzymes (Figure 2). Gene products for 33 out of the 168 PCWDN genes have been biochemically characterized in *N. crassa* (20-24,46-50,58-68) (Table S7). An additional extensive literature search was performed to determine if ortholog/paralogs of *N. crassa* PCWDN genes had been biochemically characterized in other filamentous fungi (39,69,70) (Table S7). We used OrthoMCL (71,72) to determine the orthology/paralogy of PCWDN genes in other filamentous fungi (Methods). In this way, biochemical-based annotation support was obtained for 113 out of the 168 PCWDN genes (Table S7).

### Feature Matrix and Annotation Confidence Score

The combined annotation information for PCWDN genes from the five different data types was captured in a feature matrix (Figure 3; Table S1). We next devised a simple method based on the feature matrix to assign annotation confidence scores to PCWDN genes. A differential weighting system was used to account for the different levels of confidence associated with the annotation information for each of the five heterogeneous data types with each annotation level superseded by the next (Figure 3). Annotation support from biochemical characterizations was given the highest level of confidence (factor of 16), followed by published mutant phenotypes (factor of 8), proteomic data (factor of 4), transcriptomic data (factor of 2), and functional genomics-based predictions (factor of 1). To obtain an overall annotation confidence score in the range of 0 to 1, the weighted sum of evidence support from the five data types was normalized by dividing by the maximal possible score of 31 (Figure 3). Note that the chosen factors in the differential weighting system are the simplest possible that enable mathematical mapping from diverse evidence support values to a unique annotation confidence score. Moreover, the chosen factors are such that the contribution from a given data type to the confidence score is always more than the combined contribution from all other data types with less confidence. That is, a gene with annotation support from only biochemical characterization has a higher confidence score than a gene with combined annotation support from genetics, proteomics, transcriptomics and functional genomics-based information. Table S1 lists the annotation confidence scores for the 168 PCWDN genes, and Figure 3 gives the confidence scores for the genes belonging to CAZY class AA9 (Auxiliary Activity Family 9) encoding LPMOs involved in cellulose degradation (44,47,50). For this particular case, the majority of genes encoding AA9 LPMOs within class 1 (3 out of 5 genes) and class 2 (2 out of 3 genes) are well characterized, while this is true for only 1 out of 6 genes encoding class 3 AA9 LPMOs.

### Comparative Global Transcriptome Analysis

To better define the genome-wide response of *N. crassa* to plant cell wall substrates, we analyzed combined RNA-seq data for possible correlations between the response patterns for various carbon sources. Figure 4A shows the level of correlation across the whole transcriptome for each pair of growth conditions. Hierarchical clustering of the pairwise correlation matrix across carbon sources revealed four main clusters. Avicel (cellulose) and MLG (a hemicellulose component) were in the first cluster with highly correlated gene expression patterns, likely due to the fact that both polysaccharides have a backbone that is rich in β-1,4-linked D-glucose residues. Similarly, sucrose and starch conditions fell into a second cluster, while xylan and pectin conditions were in a third cluster. This analysis also identified a fourth cluster where the gene expression in two hemicellulose components, mannan and XG, were very similar to the NoC starvation condition. In support of these observations, the *N. crassa* WT strain was observed to grow poorly on both mannan and XG. Principal component analysis of expression data under the different conditions gave a similar result (Figure 4B), where the first two principal components together explained more than 79% of the total variance.

**Figure 4:**
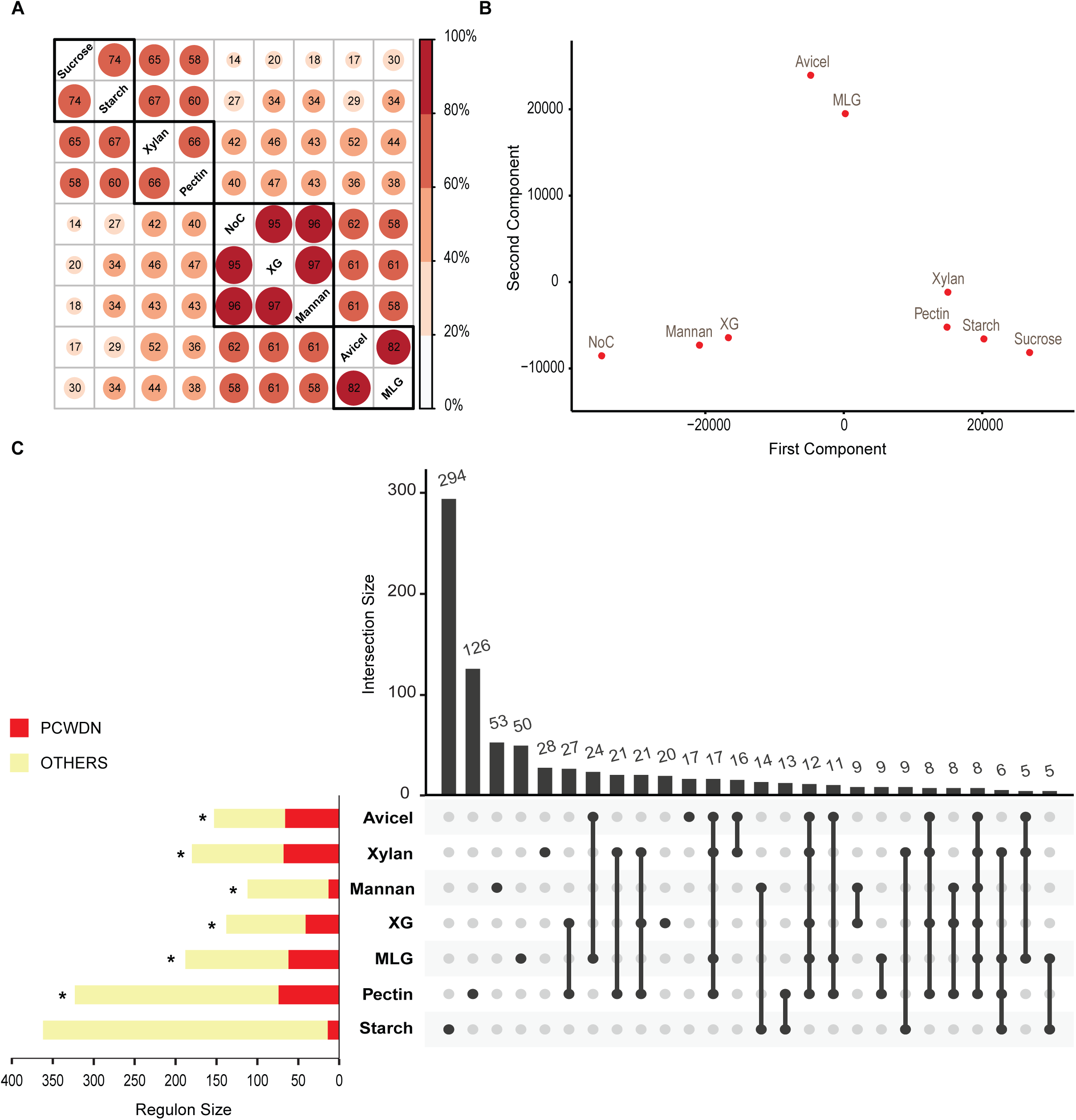
Comparative analysis of global transcriptome profiles of *N. crassa* in nine conditions. (**a)** Correlation of the whole transcriptomes across each pair of conditions. Hierarchical clustering of the pairwise correlation matrix led to the identification of four clusters of highly correlated conditions. **(b)** Principal component analysis of the whole transcriptomes in all nine conditions. The first and second principal components explain greater than 58% and 20%, respectively, of the total variance. Thus, the first two components together explain more than 79% of the total variance. **(c)** Horizontal bar plot showing the sizes of regulons for seven conditions and the overlap of each regulon with the 168 PCWDN genes. Statistically significant overlaps (p < 10^−6^) were marked with an asterisk. The vertical bar plot shows the 26 intersection sets among the seven regulons with five or more genes and was generated using UpSetR (92).

For a more detailed comparison of the *N. crassa* transcriptional response upon exposure to different plant cell wall polysaccharides, we identified the regulons for each polysaccharide using the NoC and sucrose conditions as controls. Following Benz *et al.* (19), the regulon (or up-regulon) for a given growth condition was defined as genes that were upregulated and differentially expressed in relation to both controls. Based on this definition, the regulons for Avicel, xylan, XG, mannan, MLG, pectin and starch, were determined to contain 153, 180, 138, 112, 188, 323 and 363 genes, respectively (horizontal bar plot in Figure 4C; Table S4; for the corresponding down-regulons, see Figure S1). Although starch was determined to have the largest regulon with 363 genes, it was fairly distinct among the regulons with 294 unique genes (81%). By comparison, mannan was 47% unique, followed by pectin (39%), MLG (28%), xylan (16%), XG (15%), and Avicel (11%). The overlap between each of the seven regulons was determined and Figure 4C (vertical plot) shows all regulon comparisons with an overlap of five or more genes. Interestingly, XG and pectin regulons contained the highest overlap with 27 genes in common, followed by the Avicel regulon that overlapped most highly with MLG (24 genes), xylan with pectin (21 genes), and mannan with starch (14 genes). All 26 intersection sets from this analysis were subjected to functional category analysis based on gene annotations in FunCatDB (73) (Figure S2). Most of the intersection sets involving either Avicel, xylan, XG, mannan, MLG or pectin regulons were enriched in metabolic genes (“Metabolism and Energy”).

We next computed the relative abundance of the 168 PCWDN genes within each of the seven polysaccharide regulons and the probability of their enrichment. For Avicel, 66 PCWDN genes were represented in the 153 gene regulon, a 43% enrichment (p <10^−78^). The xylan regulon contained 68 PCWDN genes (37% enrichment; p < 10^−76^), the mannan regulon contained 13 PCWDN genes (29%; p < 10^−7^), the XG regulon contained 41 PCWDN genes (43%; p < 10^−39^), the MLG regulon contained 62 PCDWN genes (32%; p < 10^−64^), the pectin regulon contained 74 PCWDN genes (22%; p < 10^−65^), and the starch regulon contained 14 PCWDN genes (3.8%; p < 10^−2^) (Figure 4C). The observed overlap between PCWDN genes and regulons was statistically highly significant, since the PCWDN genes account for only 1.7% of genes (168 out of 9758) within the *N. crassa* genome. Thus, our PCWDN genes capture a substantial (and relevant) part of the transcriptional response of *N. crassa* upon exposure to Avicel (cellulose), xylan, mannan, XG, MLG and pectin.

### Clustering of Transcriptome Profiles within the Context of the PCWDN

Comparative global transcriptional analyses underlined the importance of PCWDN genes for adaptation to different plant cell wall polysaccharides. We next performed hierarchical clustering (74) of the RNA-seq data for the 168 PCWDN genes in all nine conditions (Avicel, xylan, XG, mannan, MLG, pectin, starch, sucrose and NoC). The conditions (x-axis in Figure 5) clustered similarly to what was obtained from correlation and principal component analysis of the genome-wide transcriptome profiles presented in Figure 4A and B. The genes (y-axis in Figure 5) were grouped into six major clusters (Methods; Table S8). The 33 genes in the largest cluster were highly expressed under Avicel and MLG as compared to other conditions. Notably, this cluster contains most of the endo-β-1,4-glucanases, exo-β-1,4-glucanases, β-glucosidases, AA9 LPMOs and the identified cellodextrin transporters. The second cluster contained 15 genes and showed highest expression on xylan and consisted of genes encoding endo-xylanases, β-xylosidases, acetyl xylan esterases, xylose reductase, xylulokinase and xylitol dehydrogenase. The 10 genes in the third cluster were most strongly expressed on pectin and included genes encoding pectin methyl esterases, a rhamnogalacturonan lyase, a rhamnogalacturonan acetyl esterase, an endo-β-1,4-galactanase and a β-galactosidase. The fourth cluster composed of 27 genes showed transcriptional activity on both xylan and pectin and consisted mainly of genes encoding β-xylosidases, exo- and endo-arabinanases, α-arabinosidases, β-galactosidases, and genes involved in L-rhamnose and D-galacturonic acid metabolism. The fifth cluster composed of 6 genes with broader expression pattern contained four sugar transporters, and the final cluster composed of 29 genes contained those factors most strongly induced during starvation (excluding mannan and XG).

**Figure 5:**
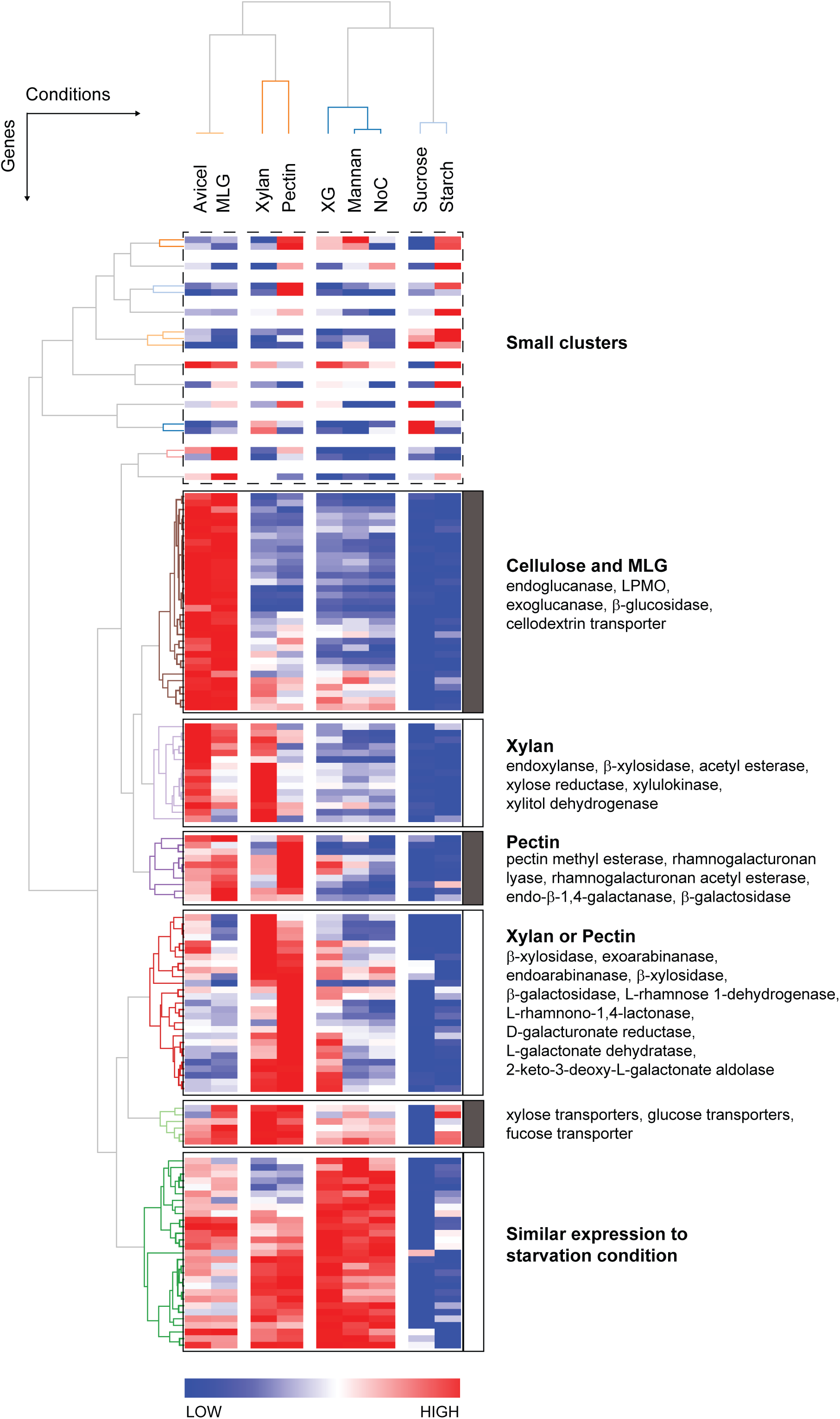
Hierarchical clustering of transcriptome profiles within the context of the PCWDN of *N. crassa*. The heatmap shows the result of two-dimensional clustering of the RNA-seq data for all 168 PCWDN genes in nine conditions corresponding to Avicel, xylan, XG, mannan, MLG, pectin, starch, sucrose and NoC, which led to the identification of six major clusters. Note that the clustering of conditions was similar to that obtained from correlation and principal component analysis of the global transcriptome profiles (Figure 4A,B).

The AA9 LPMOs, previously annotated as CAZY class GH61, degrade cellulose by oxidative cleavage (44,46,47,49). Most AA9 LMPOs were highly expressed on Avicel and MLG and their expression was correlated across the two conditions (Figure S3). In comparison to Avicel and MLG, the expression of 10 of the 14 LPMOs was much lower on XG and negligible on starch. Based on these observations, we concluded that these genes are specifically induced by non-substituted β-D-glucans and hypothesize that LPMOs in AA9 CAZY class also act on MLG, which is now included in the PCWDN of *N. crassa* (Table S1).

A recent study (49) suggested that the product of AA9 LPMO gene NCU02916 can also act on XG. However, NCU02916 was not induced when grown on XG, and the *N. crassa* WT strain (FGSC 2489) grew poorly on XG, with the transcriptome profile highly correlated with that of the NoC control (Figure 4A). Thus, it is difficult to rule out the possibility that the AA9 LPMOs also act on XG, and we therefore included such reactions in the PCWDN of *N. crassa* (Table S1). However, another recent study (50) characterized a starch-specific LPMO, NCU08746, and this gene is classified in the new CAZY class AA13 (41). Consistent with this finding, NCU08746 was differentially upregulated under starch conditions (Table S4); NCU08746 was assigned to starch-specific LPMO reaction in the PCWDN (Table S1).

### Regulation of the PCWDN by Key Transcription Factors

Previous research (16,17) on plant cell wall deconstruction by *N. crassa* led to the identification of two essential transcription factors (TFs), CLR-1 (NCU07705) and CLR-2 (NCU08042), for cellulose utilization, and one essential TF, XLR-1 (NCU06971), for utilization of xylan. All three TFs are conserved across ascomycete fungi (16,75-80). Using next generation sequencing of chromatin-immunoprecipitated DNA (ChIP-seq), a recent study (81) identified binding regions for CLR-1, CLR-2 and XLR-1 across the *N. crassa* genome under sucrose, Avicel, xylan and NoC conditions. CLR-1, CLR-2 and XLR-1 bound to regulatory regions of 293, 164 and 84 genes, respectively, in the *N. crassa* genome (81). Integrating this information, we determined that CLR-1, CLR-2 and XLR-1 bound promoter regions of 27 (p < 10^−12^), 37 (p < 10^−30^) and 20 (p < 10^−17^) genes, respectively, within the 168 PCWDN genes (Figure 6). Of the 27 PCWDN genes regulated by CLR-1, 21 are involved in cellulose utilization, including endo-β-1,4-glucanases, AA9 LPMOs, exo-β-1,4-glucanases, β-glucosidases or cellodextrin transporters, while 17 out of 20 PCWDN genes regulated by XLR-1 are involved in xylan utilization including endo-β-1,4-xylanases, β-xylosidases, α-arabinofuranosidases or xylodextrin transporters (81) (Figure 6). However, in the case of CLR-2, only 23 out of 37 PCWDN genes directly regulated by the TF are involved in cellulose utilization, while 12 other PCWDN genes are involved in xylan or mannan utilization (81) (Figure 6). These observations suggest that CLR-2, unlike CLR-1 or XLR-1, has a broader regulatory role in plant cell wall deconstruction, and which is not limited to cellulose utilization.

**Figure 6:**
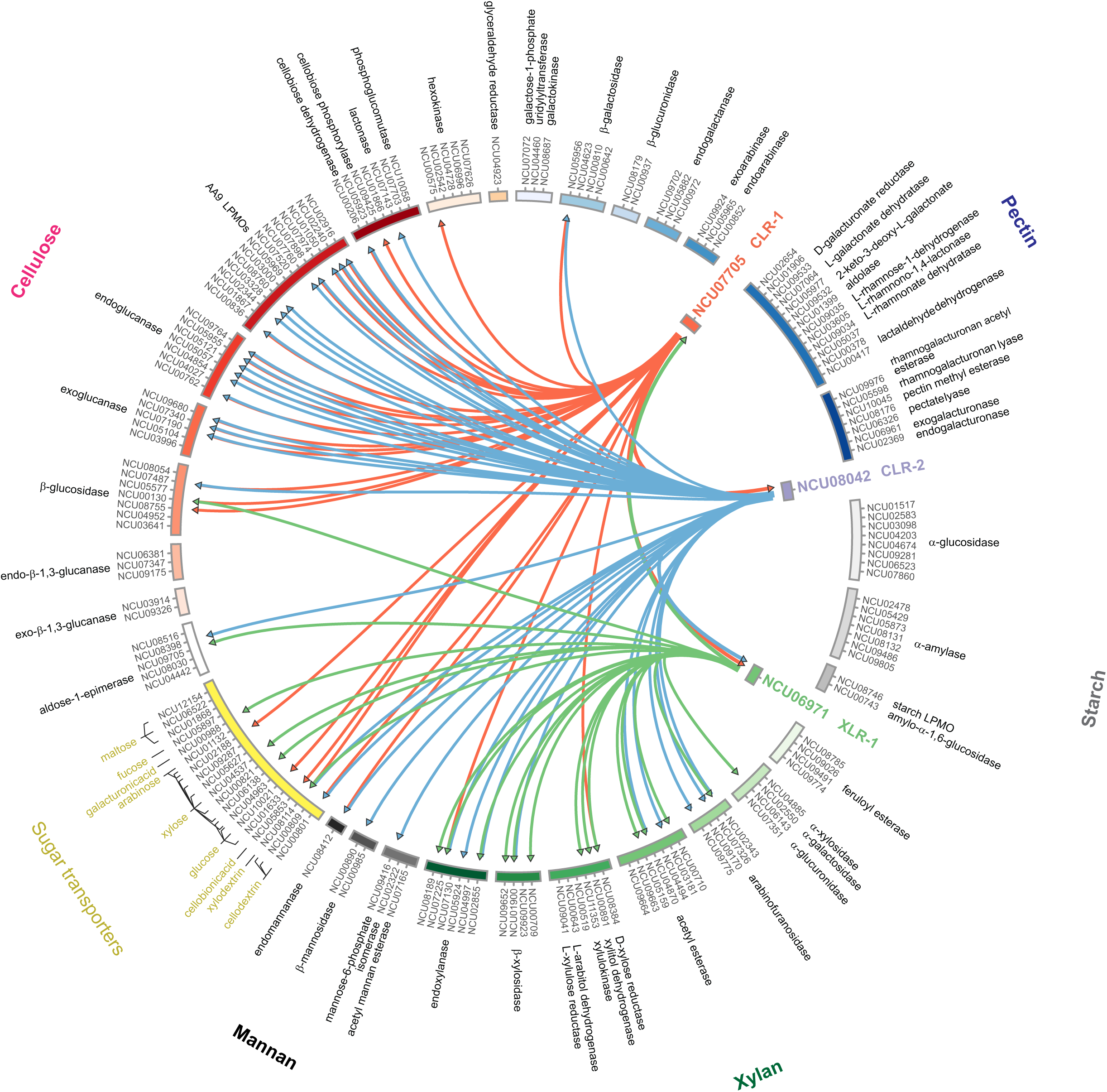
Direct regulation of the PCWDN genes by key transcription factors for cellulose and xylan utilization. The PCWDN genes have been grouped based on their biochemical function and participation in degradation pathways of different plant cell wall polysaccharides. It is seen that CLR-1 directly regulates mostly genes involved in cellulose utilization, XLR-1 directly regulates mostly genes involved in xylan utilization, while CLR-2 has a much broader role in the regulation of genes involved in plant cell wall deconstruction.

CLR-1 is known to directly regulate *clr-2* (81). *Intriguingly, we found that 19 out of 27 PCWDN genes directly regulated by CLR-1 are also directly regulated by CLR-2 (Figure 6). These observations indicate that CLR-1 and CLR-2 predominantly employ feed-forward loops (FFLs) (82-86) to regulate the cellulose utilization pathway within the PCWDN of N. crassa*.

### *clr-2, clr-1* and *gh5-7* are Important for Mannan Degradation

Our comparative transcriptome analysis revealed that the pattern of gene expression of *N. crassa* WT exposed to mannan was highly correlated to the NoC starvation condition. In contrast to the cellulose and xylan utilization pathways, the *N. crassa* genome has a mostly non-redundant mannan utilization pathway, with just one predicted endomannanase, *gh5-7* (NCU08412), and one intracellular β-mannosidase, *gh2-1* (NCU00890) (Figure 7A; Table S1). Based on ChIP-seq data (81), CLR-2 directly regulates both genes, while CLR-1 directly regulates only *gh5-7* and XLR-1 does not directly regulate any genes in the mannan pathway (Figure 6). Additionally, *gh5-7* and *gh2-1* were induced on mannan, albeit to a much lower extent than on Avicel or MLG, a context where these enzymes should not be directly required (Figure 7B). By contrast, the *clr-1* and *clr-2* genes were highly expressed in Avicel and MLG conditions, but not induced by mannan (Figure 7B). These observations led us to hypothesize that CLR-1 and CLR-2 may be important for the induction of *gh5-7,* constituting the first step in the mannan utilization pathway.

**Figure 7:**
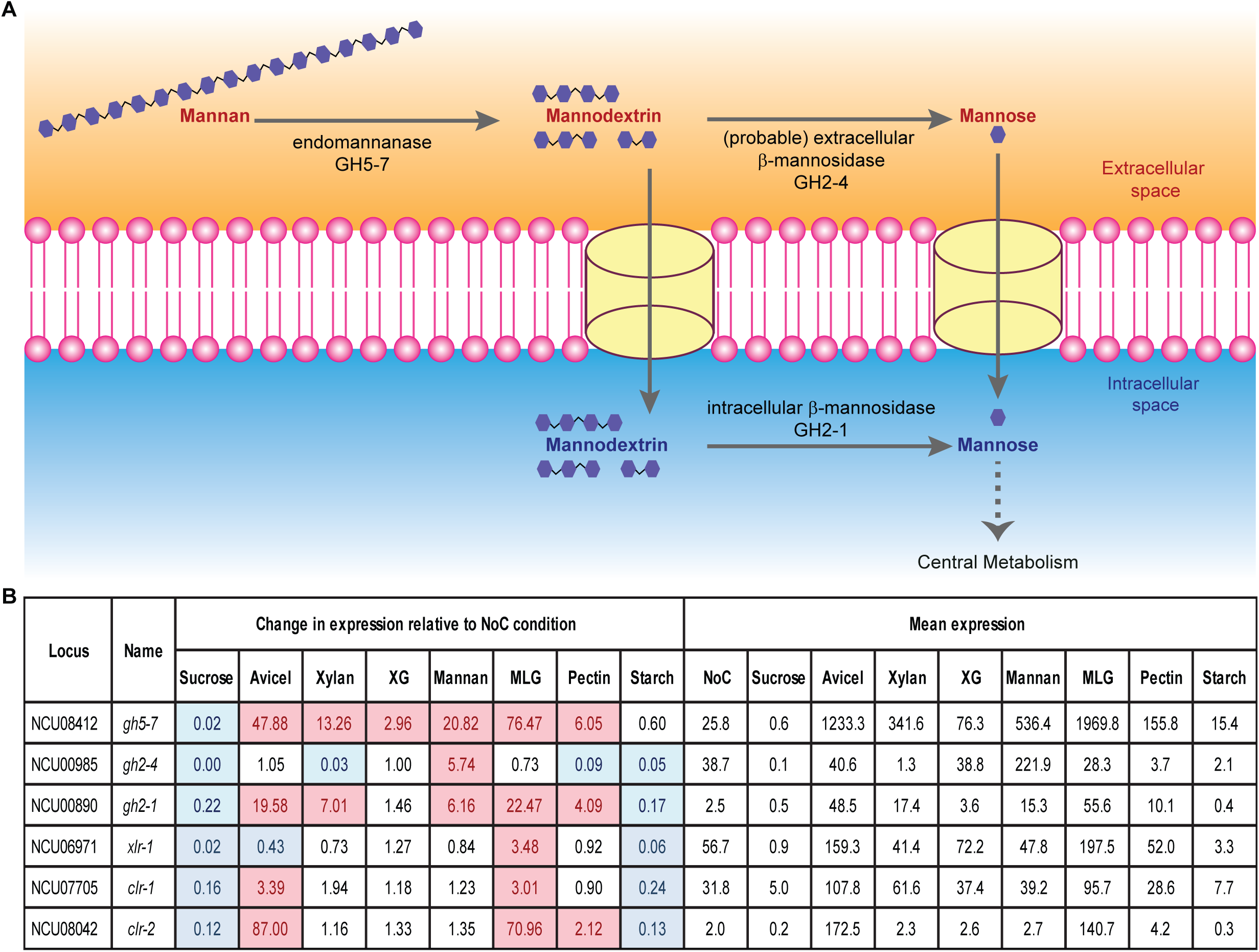
**(a)** Schematic diagram of the mannan degradation pathway. **(b)** Expression of genes encoding enzymes in the mannan degradation pathway and key transcription factors for cellulose and hemicellulose utilization across different conditions. Expression values for genes with more than 2-fold up-regulation relative to the starvation condition (NoC) are shaded in pink, while those with more than 2-fold down-regulation are shaded in blue. XG: xyloglucan; MLG: mixed-linkage glucan.

To test this hypothesis, we first designed experiments to confirm that GH5-7, as the only predicted endomannanase in *N. crassa*, is critical for mannan utilization. As the deletion strain for *gh5-7* was not available, we constructed a Δ*gh5-7* strain (ΔNCU08412) following standard procedures ((87); Methods). Since pure mannan acts as a poor inducer (Figure 8D), we used Konjac glucomannan instead of pure mannan to test our hypothesis. As predicted, the Δ*gh5-7* mutant was found to have a strong growth phenotype, accumulating only 50-60% of WT biomass when grown on glucomannan as a sole carbon source (Figure 8A, B). Additionally, both the Δ*clr-1* and Δ*clr-2* strains also grew poorly on glucomannan, accumulating only 10-20% of WT biomass, while the Δ*xlr-1* strain showed no significant growth phenotype (Figure 8A, B).

**Figure 8:**
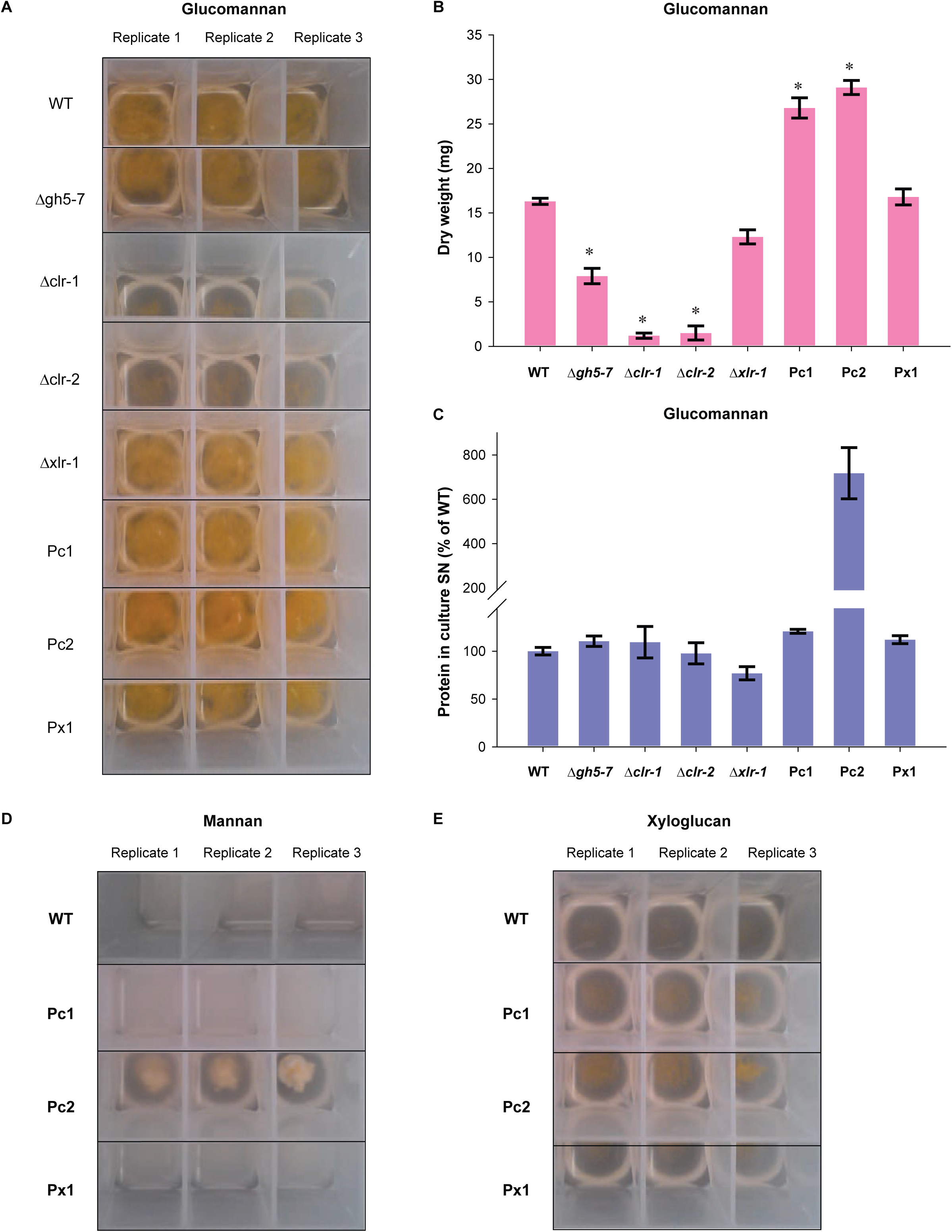
*clr-2* plays a major role in mannan and xyloglucan (XG) degradation. **(a-c)** Growth phenotypes of WT, Δ*gh5-7*, Δ*clr-1*, Δ*clr-2*, Δ*xlr-1*, Pc1 (P*ccg-1-clr-1*), Pc2 (P*ccg-1-clr-2*) and Px1 (P*ccg-1-xlr-*1) strains of *N. crassa* in medium containing glucomannan as the sole carbon source after growth for 4 days. **(a)** Photograph of 3 ml cultures with replicates in 24-well deep-well plates. **(b)** Fungal dry weights after 4 days. Bars represent standard deviations. The asterisk indicates a significant difference from WT with an unadjusted *P*-value of < 0.003 using one-way Anova. **(c)** Secreted protein in culture supernatants (SN) relative to WT. Bars represent standard deviations. The concentration of secreted protein is shown relative to WT, which was set to 100%. **(d)** Growth phenotypes of WT, Pc1, Pc2 and Px1 strains of *N. crassa* in medium containing pure mannan as the sole carbon source after growth for 4 days. **(e)** Growth phenotypes of WT, Pc1, Pc2 and Px1 strains of *N. crassa* in medium containing XG as the sole carbon source after growth for 4 days.

The results presented above showed that *clr-1*, *clr-2* and the endomannase gene *gh5-7* are important for mannan utilization by *N. crassa*. Together with the observation that CLR-2 directly regulates *gh5-7* and *gh2-1*, and that the expression of *clr-2* was negligible in the WT strain under pure mannan conditions (Figure 7B), we hypothesized that an engineered strain mis-expressing *clr-2* would show enhanced growth on mannan. A *clr-2* mis-expression strain (Pc2) constructed in a previous study (76) results in a strain that constitutively expresses *clr-2* and cellulases. We evaluated the growth of the Pc2 strain on glucomannan as compared to the WT strain and determined that the Pc2 strain accumulated 90-100% more fungal biomass than WT on glucomannan, while secreting 7x as much protein (Figure 8A-C). Similarly, a *clr-1* mis-expression strain (Pc1) accumulated 90-100% more fungal biomass than WT on glucomannan, albeit exhibiting a similar protein secretion level (Figure 8A-C). In contrast, a *xlr-1* mis-expression strain (Px1) accumulated fungal biomass at a similar level to WT on glucomannan (Figure 8A-C).

We next used Pc2, Pc1 and Px1 strains to test their growth on pure mannan as a sole carbon source. Remarkably, the Pc2 strain exhibited a robust growth phenotype on pure mannan (Figure 8D) and secreted 14x more protein than the WT strain. However, both Pc1 and Px1 strains showed a poor growth phenotype similar to WT in pure mannan (Figure 8D). Thus, these experimental results validated our hypothesis that mis-expression of *clr-2,* and thus induction of *gh5-7* and *gh2-1,* would restore growth on mannan. Furthermore, the Pc2 strain also exhibited robust growth on XG, while the Pc1 and Px1 strains showed growth characteristics that were similar to WT on XG as sole carbon source (Figure 8E). These novel insights into the regulation of the mannan and XG degradation pathway by CLR-2 will aid future efforts to engineer improved strains for degradation of lignocellulosic biomass.

### Comparison of the PCWDN with Genome-Scale Metabolic Networks of Other Fungi

A genome-scale metabolic model (iJDZ836) containing 836 metabolic genes that encode 1027 unique enzymatic activities was previously published for *N. crassa* (34) and captures biochemical reactions for catabolism of simple nutrients, central and energy metabolism and biosynthesis of biomass precursors. By comparing reactions and genes in our PCWDN with those in iJDZ836, we found that 167 out of 202 PCWDN reactions (> 82%) and 105 out of 168 PCWDN genes (> 62%) were not contained in the model (Table S9). Additionally, among the 63 common genes, 23 PCWDN genes (> 36%) had incorrect or outdated annotations (Table S9).

Apart from *N. crassa,* genome-scale metabolic models have also been reconstructed for a few other ascomycetes species, including *Aspergillus nidulans* (29), *Aspergillus niger* (30), *Aspergillus oryzae* (31), *Aspergillus terreus* (32) and *Penicillium chrysogenum* (33). Using OrthoMCL (71,72), we searched the genomes of these species for ortholog/paralogs of the 168 *N. crassa* PCWDN genes, and found 178 (*A. nidulans*), 160 (*A. niger*), 197 (*A. oryzae*), 192 (*A. terreus*) and 164 (*P. chrysogenum*) orthologous/paralogous genes (Methods; Table S9). To assess the coverage of polysaccharide degradation pathways in the reconstructed metabolic models for these species (29-33), the overlap between genes in other fungal metabolic models and ortholog/paralogs of the PCWDN genes was determined. Of the PCWDN ortholog/paralogs, 40-70% were not accounted for in the other filamentous fungal metabolic models (Table S9). These analyses highlight the significant knowledge gaps specific to pathways for degradation and utilization of plant cell wall polysaccharides in published genome-scale metabolic models for other ascomycete species.

## Conclusions

Here, we have taken a comprehensive approach, using diverse datasets, to define genes in filamentous fungi involved in the deconstruction of plant biomass, using *N. crassa* as a model (Figure 9). From these analyses, we developed hypotheses regarding regulation of mannan degradation in *N. crassa* and experimentally tested the hypothesis that the transcription factor CLR-2, previously characterized to regulate cellulose utilization (16,17), also plays a role in mannan degradation. Interestingly, *clr-2* is the ortholog of the transcriptional regulator ManR, which regulates mannan utilization in *A. oryzae* (88). These data support the view that the role of *clr-2* orthologs in the regulation of genes involved in mannan utilization is conserved among filamentous fungi. Current metabolic models for *N. crassa* (34) and other filamentous fungi (29-33) have significant knowledge gaps in the degradation pathways for most plant cell wall polysaccharides. For example, the *N. crassa* metabolic model iJDZ836 (34) has significant knowledge gaps regarding the pathways for degradation and utilization of plant cell wall polysaccharides. The iJDZ836 model contains neither the pectin degradation nor the D-galacturonic acid utilization pathway (Table S9) and lacks degradation pathways for most hemicellulosic polysaccharides including mannan, XG and MLG (Table S9). Additionally, degradation pathways for cellulose and xylan, were not captured in detail. For example, the AA9 LPMOs were incorrectly annotated as endo-β-1,4-glucanases, and therefore does not contain reactions describing the oxidative cleavage of cellulose by LPMOs (Table S9).

**Figure 9:**
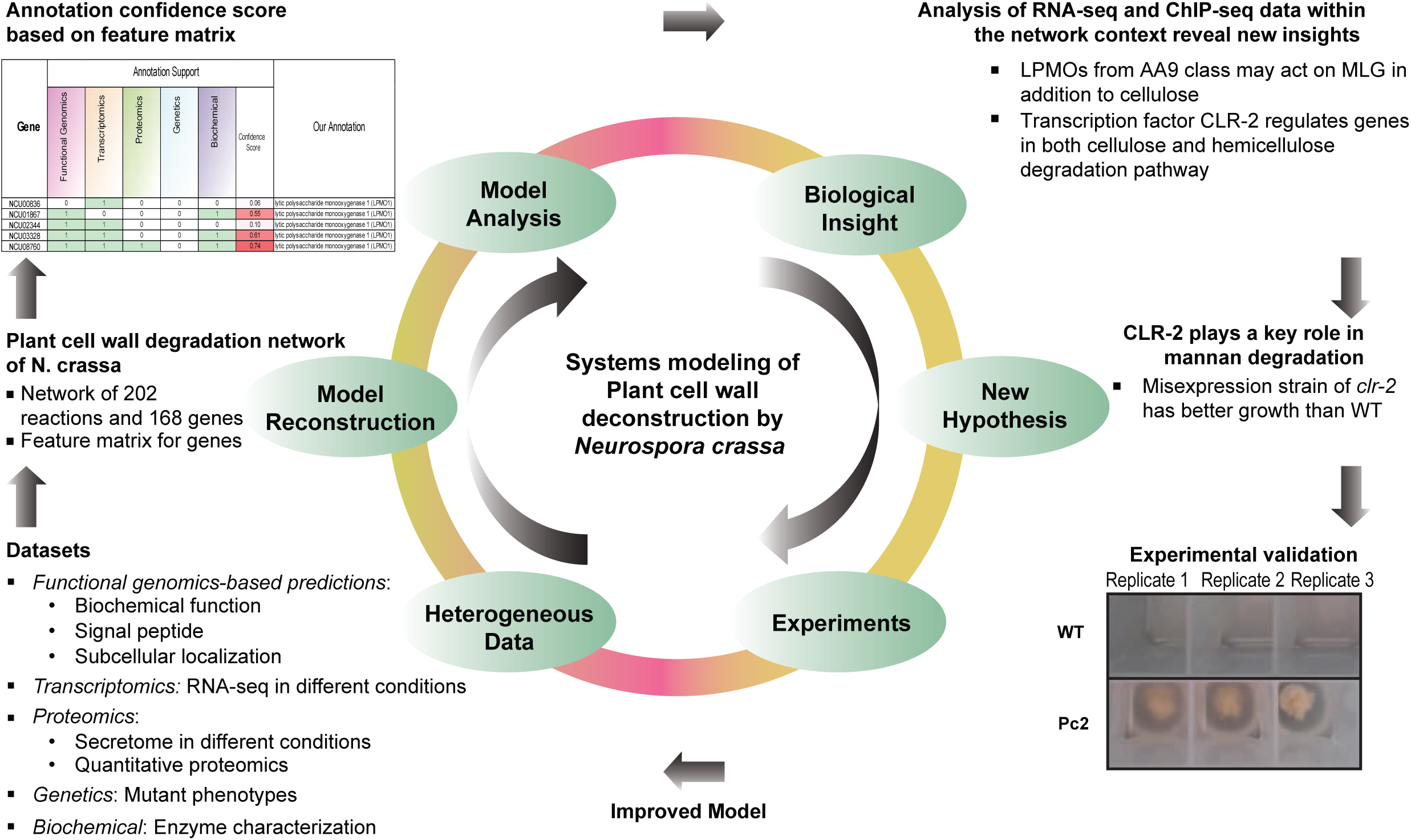
Schematic διαγραμ summarizing the systems approach undertaken here to reconstruct and analyze the plant cell wall degradation network (PCWDN) of *N. crassa.* We compiled information from diverse sources to build the network of biochemical reactions responsible for degrading plant cell wall polysaccharides into simple sugars. The combined annotation information was encapsulated in the form of a feature matrix and a simple method was devised to assign an annotation confidence score to each gene in the PCWDN. To demonstrate the utility of our PCWDN, we performed comparative transcriptomics analysis using RNA-seq data and integrated genome-wide binding data from ChIP-seq experiments for three key transcription factors regulating the plant cell wall degradation response. Integration of next generation sequencing data within the PCWDN led to the hypothesis that CLR-2 is a key TF for deconstruction of mannan and XG, a hypothesis that was subsequently validated through experimentation.

Likewise, the metabolic models for other filamentous fungi do not capture the LPMO associated reactions and pectin degradation pathway. These limitations render the available genome-scale metabolic models for filamentous fungi not well suited for the investigation of plant cell wall deconstruction, while our PCWDN will serve as a valuable resource to fill this gap. In the future, our reconstructed network is expected to play a central role in the systems analysis of complex experimental datasets, and will yield many more novel insights into plant cell wall deconstruction by filamentous fungi.

## Methods

### Databases and Functional Genomics Tools

A list of predicted carbohydrate-active enzymes in the *N. crassa* genome was compiled from two databases, the CAZY (http://www.cazy.org/; (41)) and the *N. crassa* e-Compendium (http://www.bioinf.leeds.ac.uk/∼gen6ar/newgenelist/genes/gene_list.htm; (42)). Following manual curation, we generated an updated list of predicted carbohydrate-active enzymes in *N. crassa* (Table S2). TransportDB (http://www.membranetransport.org/; (43)) was used to obtain a list of predicted transporters in the *N. crassa* genome.

Proteins destined for the secretory pathway are likely to have a signal peptide sequence in their N-terminus, and we used two prediction tools, SignalP (http://www.cbs.dtu.dk/services/SignalP/; (51)) and Phobius (http://phobius.sbc.su.se/; (52)) to predict the presence of signal peptides in amino acid sequences of PCWDN enzymes (Table S3). In order to assign the subcellular localization of enzymes, we used WoLF PSORT (http://psort.hgc.jp/; (53)) and ProtComp (http://linux1.softberry.com/berry.phtml?topic=protcomppl&group=programs&subgroup=proloc) (Table S3).

The mycoCLAP database (http://mycoclap.fungalgenomics.ca/; (70)) was used extensively to compile the list of biochemically characterized lignocellulose-active proteins of fungal origin. OrthoMCL (http://orthomcl.org/orthomcl/; (71,72)) is a tool to identify orthologous gene pairs across eukaryotic genomes. We have used OrthoMCL to determine the ortholog/paralogs of *N. crassa* PCWDN genes in more than 40 fungal genomes.

### Strains and Culture Conditions

The *N. crassa* WT reference strain and background for all mutant strains was OR74A (FGSC 2489) (12,87). The deletion strains for *clr-1* (FGSC 11029), *clr-2* (FGSC 15835) and *xlr-1* (FGSC 11066 and 11067) were obtained from the Fungal Genetics Stock Center (FGSC; http://www.fgsc.net). The mis-expression strain for *clr-1* (Pc1), *clr-2* (Pc2) and *xlr-1* (Px1) were obtained from a previous study (76). The deletion strain for *gh5-7* (NCU08412), was not available in the FGSC collection and was constructed following standard procedures (Δ*gh5-7;* ΔNCU08412) (12,87). Briefly, the 5’ upstream and 3’ downstream genomic regions surrounding NCU08412 were PCR amplified from WT genomic DNA and joined through fusion PCR with the hygromycin phosphotransferase (hph) knockout cassette (87). The resulting amplicon was transformed into FGSC 9718 (Δ*mus-51*) and selected on hygromycin slants. A homokaryotic strain was obtained through microconidia selection on water agar plates yielding the strain (Δ*gh5-7*::*hyg*^*R*^; Δ*mus-51*).

All *N. crassa* strains were pre-grown for 24 hrs on 3 mL agar slants of Vogel’s minimal media (VMM) (89) with 2% sucrose, at 30°C under dark conditions. The slants were placed under constant light at 25°C to stimulate conidia production. For flask cultures, conidia were collected and inoculated (10^6^ conidia/mL) into 100 mL liquid VMM (2% sucrose) at 25°C, under constant light and shaking (200 rpm).

### Media Shift Experiments

Media shift experiments were performed in triplicate and followed the procedure described earlier in Coradetti *et al.* (16) and Znameroski *et al.* (18) to ensure optimal comparability with the previously published RNA-seq datasets. First, using shake-flasks (200 rpm), *N. crassa* cultures were pre-grown from conidia for 16 hrs in 100 mL of VMM (89) with 2% sucrose. Next, the mycelia were passed over a Whatman glass microfiber filter and washed three times with VMM without a carbon source (NoC). The mycelial mass was then transferred to new shake flasks with 100 mL of VMM containing a specific carbon source (2% XG (P-XYGLN, Megazyme) or 2% mannan (P-MANCB, Megazyme) or 2% MLG (P-BGBM, Megazyme) or 2% Starch). After 4 hrs in the new carbon source, the mycelia were harvested over a filter, flash frozen in liquid nitrogen and stored at −80°C. Total RNA was extracted for library generation using the standard procedures as described in Tian *et al.* (15).

### RNA Sequencing and Data Analyses

Single end libraries were prepared for RNA sequencing (RNA-seq) using an Illumina kit (RS-100-0801) following standard protocols as described in Coradetti *et al.* (16). The cDNA libraries were sequenced on the Illumina HiSeq 2000 platform at the Vincent J. Coates Genomic Sequencing Laboratory, University of California, Berkeley. Three independent biological replicates of media shift cultures were sampled for *N. crassa* WT strains grown on VMM (89) with one of 4 carbon sources: XG, mannan, MLG or starch. Profiling data for each of these carbon sources is contained in Table S4 and available at the Gene Expression Omnibus database (GEO; http://www.ncbi.nlm.nih.gov/geo/; accession no. GSE90611). Previous studies (16,19) generated profiling data for *N. crassa* WT strain grown in VMM with one of 6 carbon sources (sucrose, cellobiose (CB), Avicel, xylan, pectin and OPP) or No Carbon (NoC) source, and these libraries were downloaded from the GEO database (Table S4).

Sequenced libraries were mapped to the current version of the *N. crassa* OR74A genome (v12) using Tophat v2.0.5 (http://tophat.cbcb.umd.edu/; (90, 91)). Transcript abundance was estimated in FPKM (fragments per kilobase of transcript per million mapped reads) using Cufflinks v2.0.2 (http://cufflinks.cbcb.umd.edu; (90,91)) with options of upper quartile normalization and mapping against reference isoforms. Differential expression analysis was conducted using Cuffdiff v2.0.2 (90,91). Genes with a multiple-hypothesis adjusted p-value below 0.05 and at least two-fold induction were determined to be significantly differentially expressed between conditions.

Starting from average FPKMs of genes across RNA-seq library replicates for a condition, hierarchical clustering was performed using Cluster 3.0 software suite (http://bonsai.hgc.jp/∼mdehoon/software/cluster/software.htm; (74)). Before clustering, genes were filtered out that displayed consistently low expression (<10 FPKM) in all conditions.

FPKM were log-transformed, normalized across conditions, and centered on the geometric mean across conditions on a per gene basis. The average linkage method was used for cluster generation, with Pearson’s correlation as similarity measure. Visualization of clusters was performed using GENE-E software suite (http://www.broadinstitute.org/cancer/software/GENE-E/).

### Growth Assays

Growth assays on cell wall substrates were performed in 3ml liquid cultures in 24-well plate format (GE Healthcare Life Sciences 7701-5102 with breathable sealing tape Fisher Scientific 1256705). 10^6^ conidia/mL were inoculated into VMM with 0.5% wt/vol 1,4-β-D-mannan from carob (Megazyme P-MANCB), Konjac glucomannan (Megazyme P-GLCML), or xyloglucan from tamarind (Megazyme P-XYGLN) as the carbon source.

Cultures were grown 48 hrs at 25°C with constant light and shaking at 250 rpm. At the end of the incubation, mycelia were concentrated by centrifugation at 3000 RCF for 10 minutes. Culture supernatants were then assayed for soluble protein with Bio-Rad protein assay dye reagent (Bio-Rad 500-0006), using bovine serum albumin (NEB 9001S) as the protein standard. Mycelia were washed twice in water and lyophilized before weighing for biomass determination.

## Acknowledgements

The authors thank Vincent Wu, Carly Grant and Hillary Tunggal for technical assistance, and Xin Li and William Beeson for insightful discussions. This work was supported by grants from the Energy Biosciences Institute to N.L.G. and N.D.P., and National Institutes of Health NRSA Trainee Grant 2 TR32 GM 7127-36 A1 to S.T.C. The authors also thank the FGSC and the *Neurospora* Genome Project for continuous support, and acknowledge use of materials generated by NIH grant P01 GM068087 “Functional analysis of a model filamentous fungus”.

## Authors’ contributions

A.S., J.P.C., S.T.C., J.P.B., N.D.P. and N.L.G. designed research; A.S., J.P.C., S.T.C. and J.P.B. performed research; A.S., J.P.C., S.T.C., J.P.B. and N.L.G. analyzed data; and A.S., J.P.C., J.P.B., J.A.E., N.D.P. and N.L.G. wrote the paper.

## Supplementary Figure Captions

**Figure S1:**
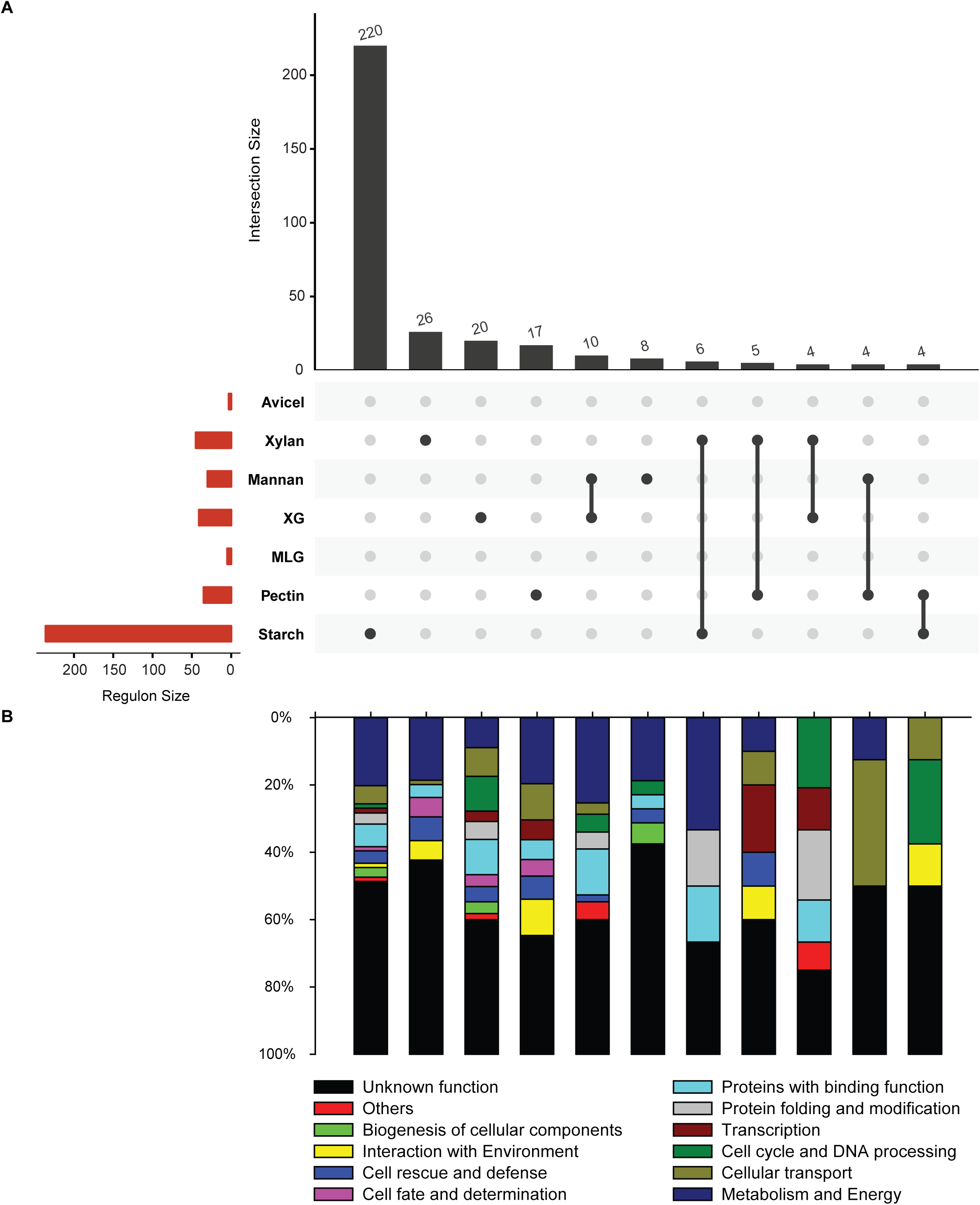
Comparative analysis of the down-regulons for the seven conditions. Analogous to up-regulon (for brevity referred to as ‘regulon’ in main text), the down-regulon for a growth condition can be defined as genes that were downregulated and differentially expressed in comparison to both NoC and sucrose controls. **(a)** Horizontal bar plot shows the size of the down-regulons for seven conditions. The down-regulons for Avicel, xylan, XG, mannan, MLG, pectin and starch, were determined to contain 3, 45, 41, 30, 5, 35 and 236 genes, respectively (Table S4). Vertical bar plot shows the 11 intersection sets among the seven down-regulons with 4 or more genes and was generated using UpSetR (92). It is seen that 220 out of 236 genes in the starch down-regulon, 26 out of 45 genes in the xylan down-regulon, 20 out of 41 genes in the XG down-regulon, 17 out of 35 genes in the pectin down-regulon, and 8 out of 30 genes in the mannan down-regulon, have no overlap with other down-regulons. It is also seen that 10 genes are common between XG and mannan down-regulons, 6 genes between xylan and starch down-regulons, 5 genes between xylan and pectin down-regulons, 4 genes between xylan and XG down-regulons, 4 genes between mannan and pectin down-regulons, and 4 genes between pectin and starch down-regulons. **(b)** Functional category analysis (73) of the 11 intersection sets among the down-regulons for seven conditions with four or more genes. Information on the functional category of *N. crassa* genes was obtained from Munich Information Center for Protein Sequence (MIPS) database (http://mips.helmholtz-muenchen.de/funcatDB/; (73)). The ‘Others’ category includes genes with functional categorization different from the 10 categories listed in the legend. The ‘Unknown function’ category includes genes with unclassified or unknown function. The relative contribution of a functional category to each set of genes is depicted with the total number of genes in each pool equal to 100%.

**Figure S2:**
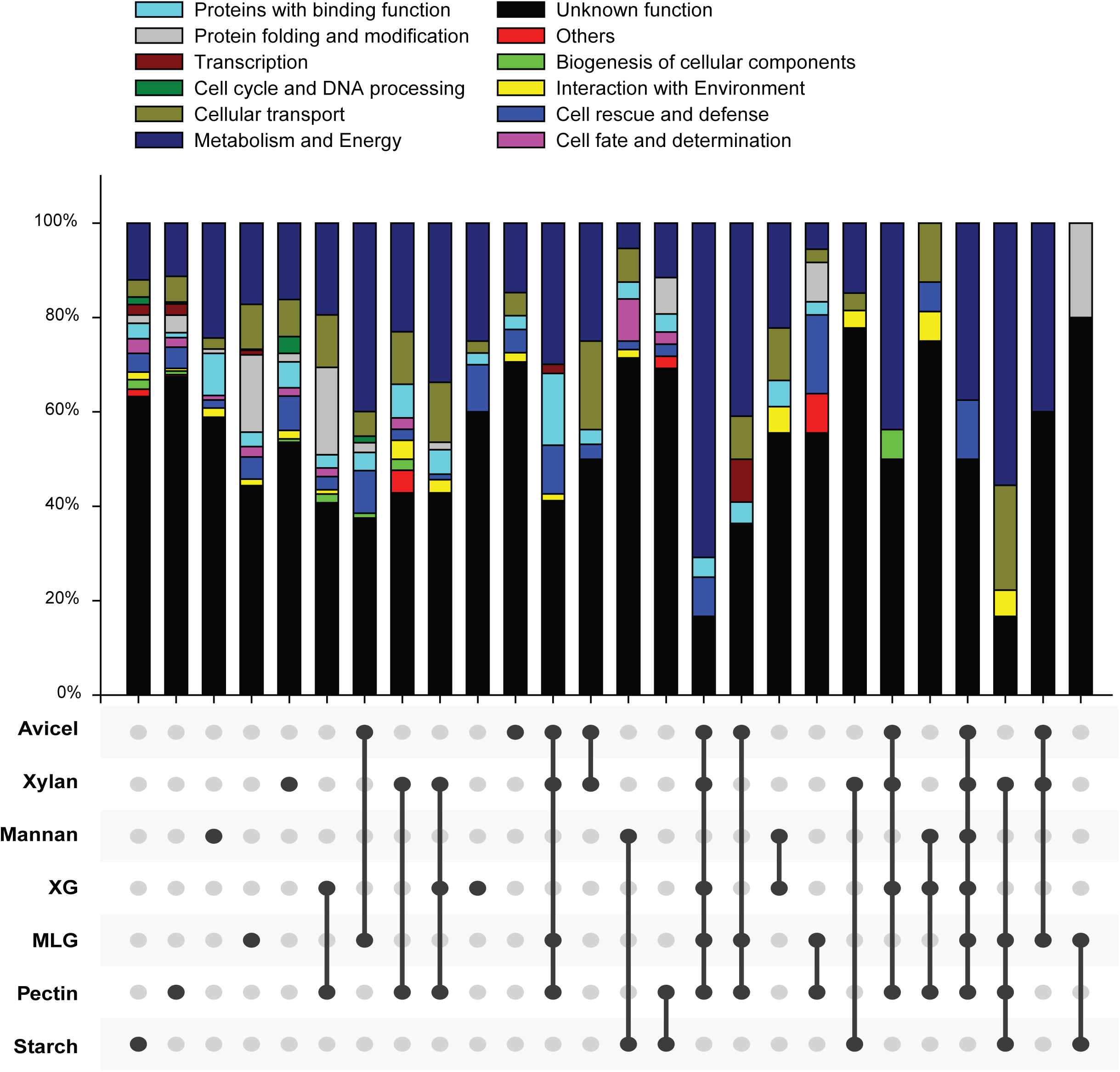
Functional category analysis of the 26 intersection sets among the up-regulons for seven conditions with five or more genes (shown in Figure 4C). Information on the functional category of *N. crassa* genes was obtained from Munich Information Center for Protein Sequence (MIPS) database (http://mips.helmholtz-muenchen.de/funcatDB/; (73)). The ‘Others’ category includes genes with functional categorization different from the 10 categories listed in the legend. The ‘Unknown function’ category includes genes with unclassified or unknown function. The relative contribution of a functional category to each set of genes is depicted with the total number of genes in each pool equal to 100%.

**Figure S3:**
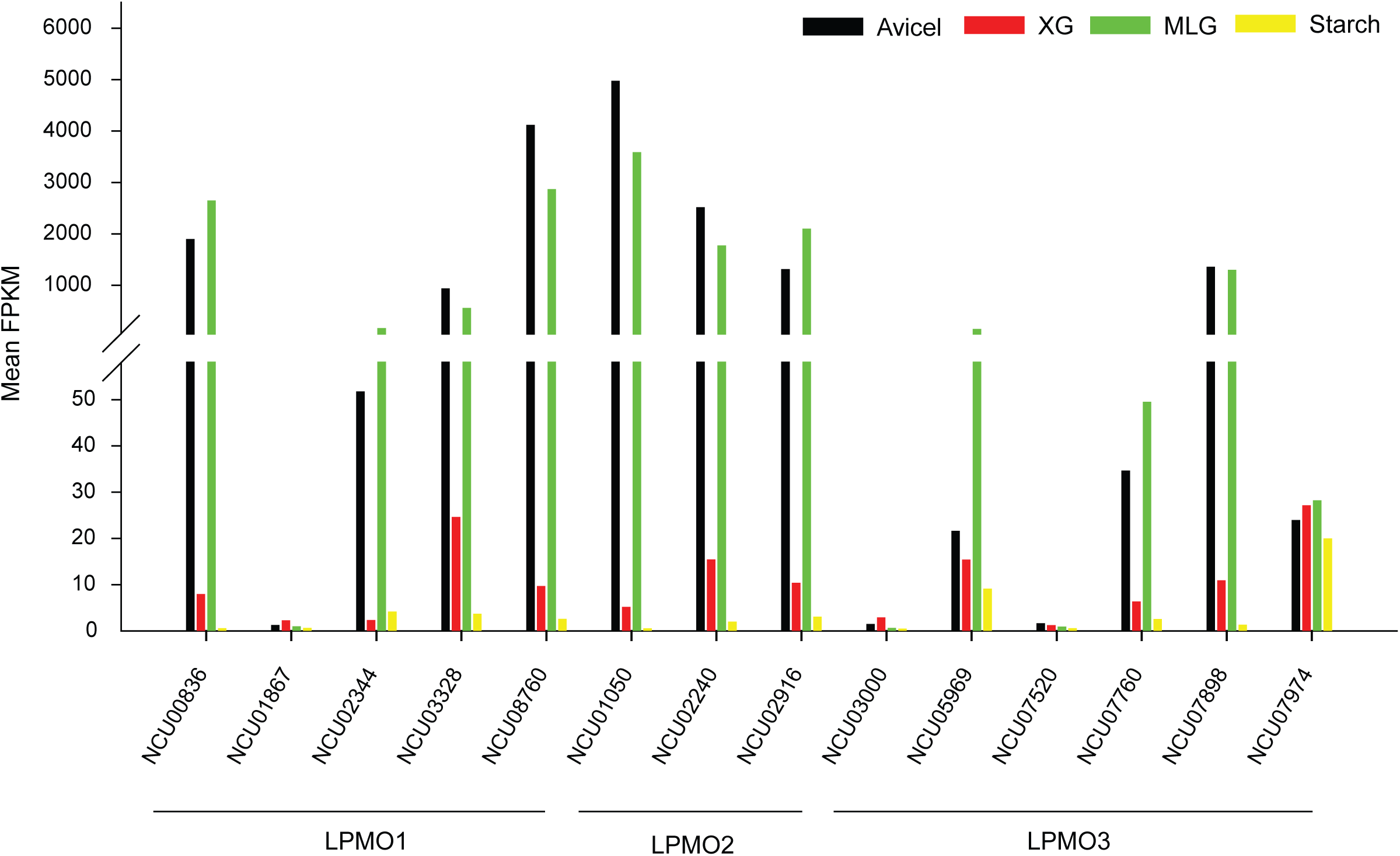
Expression of characterized and predicted AA9 LPMOs in *N. crassa* grown on four plant cell wall polysaccharides with D-glucose backbone: Avicel, xyloglucan (XG), mixed-linkage glucan (MLG) and starch. In comparison to Avicel and MLG, the expression of LPMOs was much lower on XG and negligible on starch.

### Supplementary Tables

**Table S1: List of reactions, compounds and genes in the plant cell wall degradation network (PCWDN) of *N. crassa*.** The first, second and third sheets contain the list of reactions, genes and compounds, respectively, in the PCWDN. The fourth sheet gives the participation of the PCWDN genes in the cellulose, hemicellulose, pectin and starch degradation pathways. The fifth and sixth sheets contain the feature matrix and annotation confidence scores, respectively, for genes in the PCWDN of *N. crassa* based on functional genomics, transcriptomics, proteomics and genetics data as well as biochemical characterizations. The seventh sheet contains a list of research articles utilized to reconstruct the PCWDN. The eighth sheet contains information on the structural units comprising the backbone and side chains of different plant cell wall polysaccharides such as cellulose, xylan, xyloglucan, mannan, galactomannan, glucomannan, galactoglucomannan, mixed-linkage glucan, homogalacturonan, xylogalacturonan, rhamnogalacturonan I, amylose and amylopectin. NoC: No Carbon; CB: Cellobiose; XG: Xyloglucan; MLG: Mixed-Linkage Glucan, OPP: Orange Peel Powder.

**Table S2: List of predicted genes coding for carbohydrate-active enzymes (CAZY) in *N. crassa* and the plant cell wall degradation network (PCWDN).** List of CAZY genes in *N. crassa* was obtained from two sources: CAZY database (41) and the *N. crassa* e-Compendium (42). The first sheet contains the consensus list of predicted CAZY genes in *N. crassa*. The second sheet contains the list of predicted CAZY genes in *N. crassa* from CAZY database. The third sheet contains the list of predicted glycoside hydrolase (GH) genes in *N. crassa* from e-Compendium (42) The fourth sheet contains the list of CAZY classes known to be involved in native cell wall remodelling. NoC: No Carbon; CB: Cellobiose; XG: Xyloglucan; MLG: Mixed-Linkage Glucan, OPP: Orange Peel Powder.

**Table S3: Functional genomics-based annotation support for genes in the plant cell wall degradation network (PCWDN) of *N. crassa*.** The table contains information from following sources: CAZY, BROAD, TransportDB, SignalP, Phobius, WoLF PSORT and ProtComp.

**Table S4: Transcriptomics-based annotation support for genes in the plant cell wall degradation network (PCWDN) of *N. crassa*.** The first sheet contains information on the transcriptiomics-based annotation support for each gene in the PCWDN. The second sheet contains a table with the list of RNA-seq libraries for *N. crassa* WT strain grown in different conditions along with their GEO (http://www.ncbi.nlm.nih.gov/geo/) accession numbers and references for the profiling data. The third sheet contains a table with the expression of genes in different conditions, separately, for replicate RNA-seq libraries. The fourth sheet contains a table with the average expression (Mean FPKM) of genes in each condition across replicate RNA-seq libraries. The fifth sheet contains a table with information on differential expression of genes in each condition compared to the No Carbon control. The sixth sheet contains a table with information on differential expression of genes in each condition compared to the sucrose control. The set of differentially expressed genes was computed using the Cuffdiff package (91). The seventh and eighth sheets contain genes in the up-regulons and down-regulons, respectively, of *N. crassa* in seven conditions: Avicel, Xylan, Xyloglucan, Mannan, Mixed-Linkage Glucan, Pectin and Starch. NoC: No Carbon; CB: Cellobiose; XG: Xyloglucan; MLG: Mixed-Linkage Glucan, OPP: Orange Peel Powder.

**Table S5: Proteomics-based annotation support for genes in the plant cell wall degradation network (PCWDN) of *N. crassa.*** The first sheet contains information on the proteomics-based annotation support for each gene in the PCWDN. The second sheet contains the compiled dataset of secretome for *N. crassa* strains obtained under different conditions from published literature. The third sheet contains the list of references used to compile this dataset.

**Table S6: Compiled dataset of experimentally validated deletion strains with growth-deficient phenotypes compared to WT for genes in the plant cell wall degradation network (PCWDN) of *N. crassa*.**

**Table S7: Biochemical characterization of genes in the plant cell wall degradation network (PCWDN) of *N. crassa*.**

**Table S8: Hierarchical clustering of genes in the plant cell wall degradation network (PCWDN) of *N. crassa* based on RNA-seq data obtained in nine different conditions.** The first sheet lists the different clusters in the same order as shown in Figure 5.

**Table S9: Comparative analysis of the plant cell wall degradation network (PCWDN) with the genome-scale metabolic models of *N. crassa* and other filamentous fungi.** The first and second sheets compare the list of reactions and genes, respectively, in the PCWDN and genome-scale metabolic model iJDZ836 of *N. crassa*. The third sheet gives the occurrence of orthologs or paralogs of *N. crassa* PCWDN genes in genome-scale metabolic models of other filamentous fungi.

